# EYKTHYR reveals transcriptional regulators of spatial gene programs

**DOI:** 10.1101/2025.05.19.654884

**Authors:** Spencer Krieger, Ellie Haber, Jian Ma

**Author notes:** Correspondence (J.M.).

## Abstract

Understanding how transcription factors (TFs) orchestrate gene regulatory networks that define complex tissue structures is central to uncovering tissue organization and disease mechanisms. Although spatial multiome technologies now enable *in situ* measurement of both transcriptional activity and chromatin accessibility, existing computational methods either overlook spatial tissue context or are hindered by the high dropout rates characteristic of such data. Here, we introduce Eykthyr, a computational framework that integrates gene expression and chromatin accessibility within a spatially aware model to identify TFs driving spatial gene programs. Eykthyr mitigates dropout effects by leveraging interpretable, low-dimensional embeddings of gene expression and chromatin accessibility – both linear with respect to their input – enabling robust identification and scalable inference of spatial transcriptional regulators. Applied across diverse spatial multiome datasets, Eykthyr consistently outperforms existing approaches, accurately identifying TFs that coordinate spatial gene programs in mouse brain development and regulate T-cell states within tumor microenvironments. Eykthyr establishes a foundation for decoding how TFs interpret local intercellular signaling to shape tissue structure, offering insights into the regulatory logic underlying spatial organization in health and disease.

## Introduction

Transcription factors (TFs) orchestrate spatial gene expression within complex tissues, guiding development and mediating cell-cell communication [1–3]. Acting as molecular switches, TFs regulate transcriptional programs that determine cell fate, and their disruption can lead to disease [4–6]. By interpreting extracellular cues and propagating signals through gene regulatory networks, TFs establish localized expression patterns that shape tissue organization. However, single-cell genomic technologies that lack spatial context present a major barrier to identifying TFs that govern spatial gene programs.

Recently, spatial multiome technologies have been developed to simultaneously measure transcriptional activity and chromatin accessibility *in situ* [7–9]. By correlating regulatory regions (e.g., enhancers and promoters) to their target genes, these dual-modality datasets offer new insights into TF-driven spatial gene regulation while reducing the need to infer regulatory activity solely from transcriptomic data, providing clearer potential cause-and-effect relationships in spatially resolved gene regulation. Moreover, because spatial multiome data preserve each cell’s physical location within the tissue, they offer a unique window into how local signaling, cellular neighborhoods, and microenvironmental cues influence TF-driven gene expression.

Despite these advantages, analyzing spatial multiome data remains challenging due to high dropout rates, with only 6-11% of the gene expression read count matrix typically containing nonzero values [7– 9], and even greater sparsity in chromatin accessibility data. Existing gene regulation modeling methods [10–16] are not designed for spatial multiome data; they either ignore spatial context, focus solely on cell type-level gene regulation, or struggle with extreme sparsity. As a result, no dedicated computational framework currently exists to identify TFs governing spatial gene expression patterns.

Here, we present Eykthyr, a computational method designed to identify TFs governing spatial gene programs (or metagenes) using spatial multiome data. Eykthyr addresses high dropout by generating low-dimensional embeddings of gene expression and chromatin accessibility that remain linear with respect to input, denoising the data while preserving interpretability – shifts in metagene embeddings map directly back to changes in gene expression. Eykthyr uses linear models to learn relationships between TF activity and metagene expression within each cell’s local neighborhood. These two layers of linear mappings – from TF activity to metagenes and from metagenes to gene expression – enable reasoning about how TF activity affects gene expression. We applied Eykthyr to multiple spatial multiome datasets, demonstrating its ability to identify key TFs regulating spatial gene programs in various contexts. Eykthyr’s predictions align strongly with prior knowledge, capturing TFs that influence cell fate within specific tissue regions during mouse brain development and identifying context-dependent TF effects on T cell states in distinct tumor microenvironments in human metastatic melanoma.

## Results

### Overview of Eykthyr

Eykthyr takes as input a spatial multiome dataset containing gene expression, chromatin accessibility, and spatial coordinates of each cell or spot within a tissue sample (**Fig**. 1). It aims to identify TFs that influence spatial gene programs by integrating these data modalities and analyzing region-specific interactions between TF activity and metagene expression.

**Figure 1:**
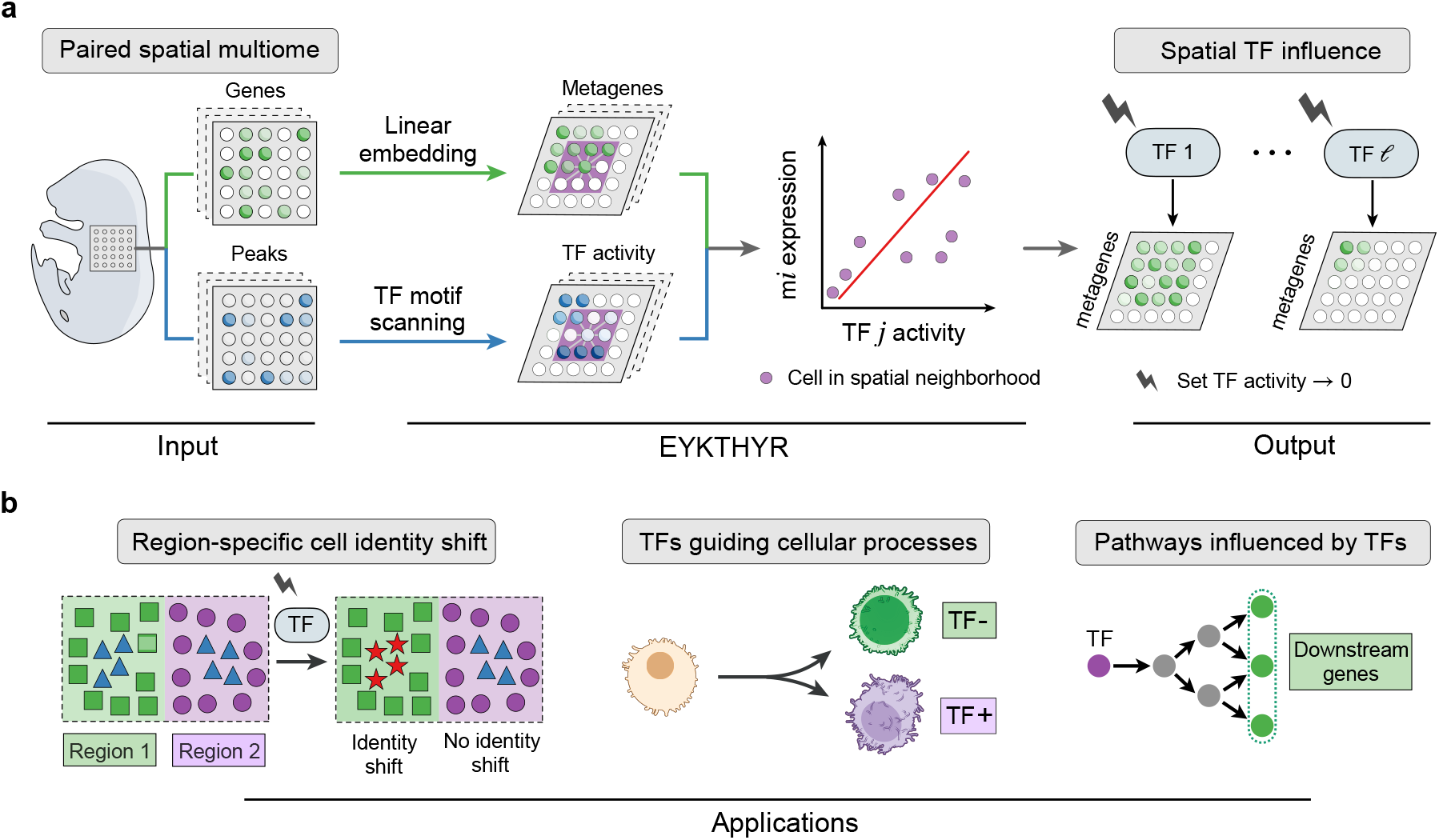
Overview of Eykthyr. **a**. Eykthyr processes paired spatial multiome data, including gene expression, chromatin accessibility, and spatial coordinates for each cell or spot. To address data sparsity, gene expression is embedded into interpretable metagenes (gene programs), while chromatin accessibility is summarized as TF activity using a TF-motif scanning approach. For each cell *c*, Eykthyr fits a regularized linear model linking the expression of metagene *m*_*i*_ with the activity of TF *j* from cells in *c*’s spatial neighborhood, estimating the interaction weight of *j* on *m*_*i*_. These weights are then used to simulate and predict changes in metagene expression following shifts in TF activity. **b**. Downstream applications of Eykthyr include predicting region-specific shifts in cell identity, identifying TFs driving cellular processes such as differentiation, and pin-pointing pathways influenced by specific TFs.

The first step in Eykthyr is generating low-dimensional embeddings for preprocessed gene expression and chromatin accessibility (**Methods**). Each embedding is linear with respect to its input, ensuring interpretability – shifts in the embedding space directly reflect changes in gene expression or chromatin accessibility. This allows us to reason about how changes in TF activity translate into shifts in gene expression patterns. For gene expression, Eykthyr employs Popari [17], a spatially aware framework of non-negative matrix factorization (NMF) [18]. Popari extends our prior work SpiceMix [19], addressing data sparsity through hierarchical binning and learning a shared set of metagenes across samples, where each metagene corresponds to a linear combination of input gene expression. This metagene embedding provides an interpretable low-dimensional representation of the gene expression landscape.

To connect chromatin accessibility with TF activity, Eykthyr estimates TF activity based on the number of accessible ATAC-seq peaks containing its binding motif. The assumption is simple but effective: the accessibility of a TF’s binding motifs across the genome correlates with its activity in the cell. This transformation provides a functional connection between chromatin accessibility and TF activity, which we leverage for downstream analysis.

Once TF activity is estimated, Eykthyr models the influence of each TF on metagene expression within the spatial context of each cell. Specifically, for each cell or spot in the tissue, Eykthyr applies regularized linear regression to infer the relationships between TF activity and metagene expression using spatially proximal cells or spots. This step effectively learns the influence of each TF on metagene expression, akin to determining edge weights in a TF – metagene regulatory network, where nodes represent TFs or metagenes and edges are directed from TFs to metagenes. By modeling this regulatory network at the metagene level rather than the individual gene level, Eykthyr effectively reduces the number of interaction weights to estimate, making it computationally feasible to infer edge weights that are distinct for each cell in the tissue.

A key strength of our method is that the linear relationships between TF activity and metagene expression allow for *in silico* predictions of how shifts in TF activity alter gene expression patterns across the tissue. This provides a framework for studying TF-driven regulation of spatial gene programs and offers insights into the regulatory mechanisms governing tissue organization.

We use these learned relationships in downstream analyses to explore how spatially localized gene programs are controlled by specific TFs. Eykthyr enables visualization of tissue-wide changes in gene expression following an *in silico* TF knockout, identification of TFs governing particular cellular processes, and discovery of pathways influenced by TF activity in specific tissue regions. These capabilities highlight the novelty of Eykthyr in linking TF activity and gene expression within a unified spatial framework.

A detailed description of Eykthyr is provided in **Methods**, and details on dataset preprocessing and hyperparameter selection are in the **Supplementary Note**.

### Decoding regulatory mechanisms in pallium differentiation

We first applied Eykthyr to examine embryonic mouse brain development using MISAR-seq data [7] (see **Fig**. 3a), focusing on progenitor cells differentiating into the ventricular region of the dorsal pallium (DPallv), the mantle region of the dorsal pallium (DPallm), and the subpallium (**Fig**. 2a). This developmental pathway serves as an ideal benchmark because it is well-characterized and involves TFs known to drive cell fate decisions. We collected a set of TFs implicated in pallium differentiation from the literature [20], including *Arx, Emx1, Emx2, Gli3, Lef1, Lhx2, Lmx1a, Nr2e1, Nr2f1, Nr2f2, Otx1, Bhlhe22, Pax6*, and *Sp8*. Verifying that Eykthyr can recover these known TFs builds confidence in its predictive capabilities.

**Figure 2:**
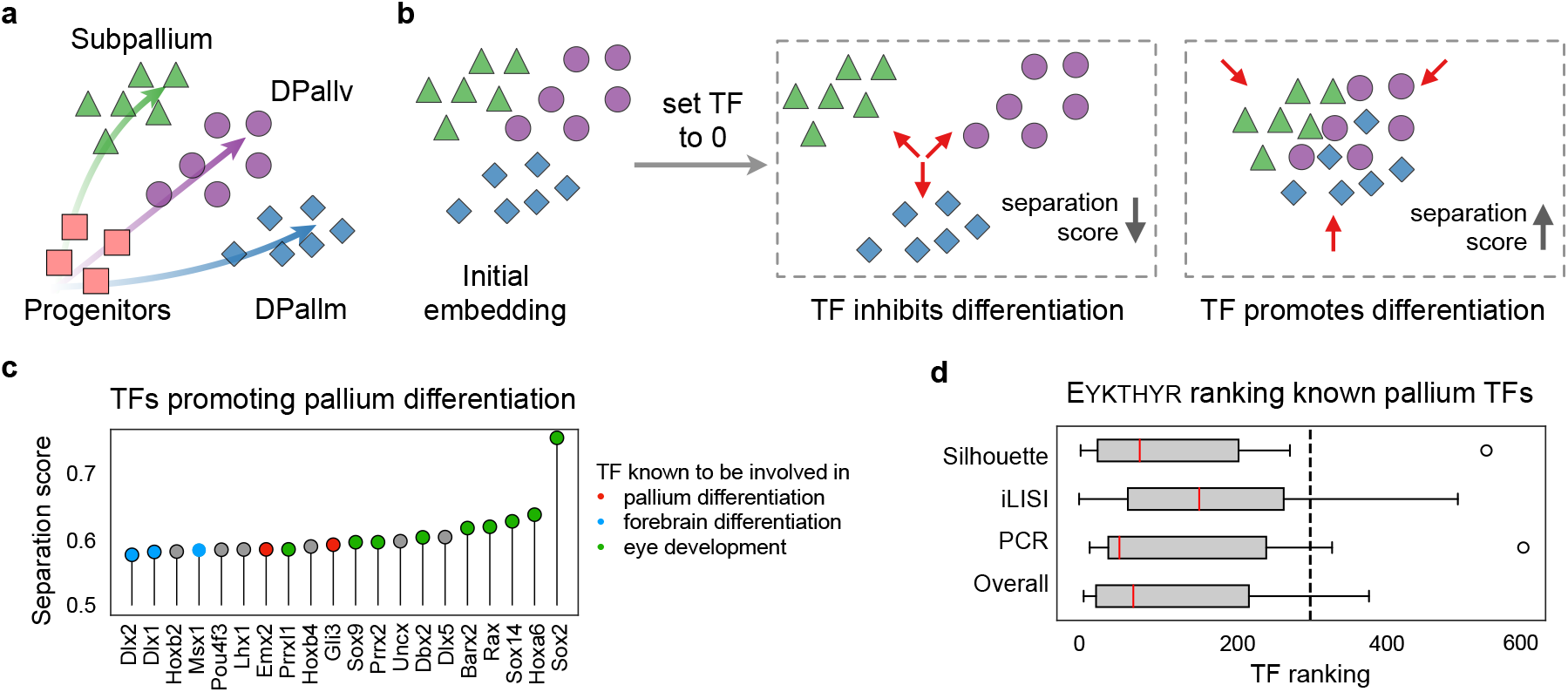
Evaluation of Eykthyr’s predictions of TFs driving pallium differentiation. **a**. Schematic illustrating the differentiation of pallium progenitors into three subtypes during mouse brain development. **b**. Measurement of each TF’s influence on pallium differentiation, determined by Eykthyr’s predicted shift in metagene embeddings after simulating TF knockout (setting its expression to zero). **c**. Top TFs identified by Eykthyr as promoting pallium differentiation, ranked using the metric in **b** (further analyzed in **Fig**. 3). **d**. Box plot showing Eykthyr’s ranking of TFs known from the literature to influence pallium differentiation. The dashed vertical line represents the expected median rank of a random set of TFs. Eykthyr consistently ranks known differentiation-promoting TFs highly, with statistically significant results.

To quantify each TF’s effect on differentiation, Eykthyr simulates a TF knockout by setting its estimated activity to zero, propagates this through learned TF-metagene weights to shift the metagene embedding, and recalculates cellular states. We then evaluate changes in cluster separation using silhouette width, iLISI, and principal component regression comparison to assess the TF’s role in maintaining distinct cell fates (**Fig**. 2b). TFs that, when knocked out, lead to collapsed differentiation clusters are inferred to be key regulators. TFs are ranked by their ability to maintain developmental separability and compared against known regulators.

Eykthyr successfully prioritized TFs involved in early brain development (**Fig**. 2c). Fourteen of the top twenty ranked TFs have known roles in pallium (*Gli3* and *Emx2*), forebrain (*Msx1, Dlx1*, and *Dlx2*), or visual cortex development (*Sox2, Hoxa6, Sox14, Rax, Barx2, Dbx2, Prrx2, Sox9*, and *Prrxl1*). TFs implicated in pallium differentiation were significantly enriched among the top-ranked TFs across all cluster separation metrics (Silhouette: *P* = 0.001; iLISI: *P* = 0.01; PCR: *P* = 0.005; combined score: *P* = 1 × 10^−4^) (**Fig**. 2d). Comparison with CellOracle [14] and an expression-only variant of Eyk-thyr showed that neither baseline prioritized these TFs better than random (**Fig**. S1a and **Fig**. S1b), highlighting Eykthyr’s ability to pinpoint spatially relevant TFs.

### *Msx1* as a central regulator in pallium development

To showcase Eykthyr’s spatially-informed predictions, we examined its analysis of *Msx1*, a TF involved in forebrain development around embryonic day 16 [21, 22]. Despite low *Msx1* transcript levels in this tissue (**Fig**. 3b, left), Eykthyr inferred strong activity in the pallium and diencephalon based on chromatin accessibility (**Fig**. 3b, right), underscoring the value of integrating accessibility data. Metagene m11, prominent in DPallv (**Fig**. 3c; other metagene expression in **Fig**. S2a), was predicted to be strongly regulated by *Msx1* (**Fig**. S2b). Transforming predicted metagene shifts into gene-level changes allowed us to rank genes likely affected by an *Msx1* knockout. These predictions were significantly enriched for validated downstream genes from an existing GRN [14] (*P* = 4 × 10^−7^; **Fig**. 3d), supporting the functional relevance of Eykthyr’s predictions.

**Figure 3:**
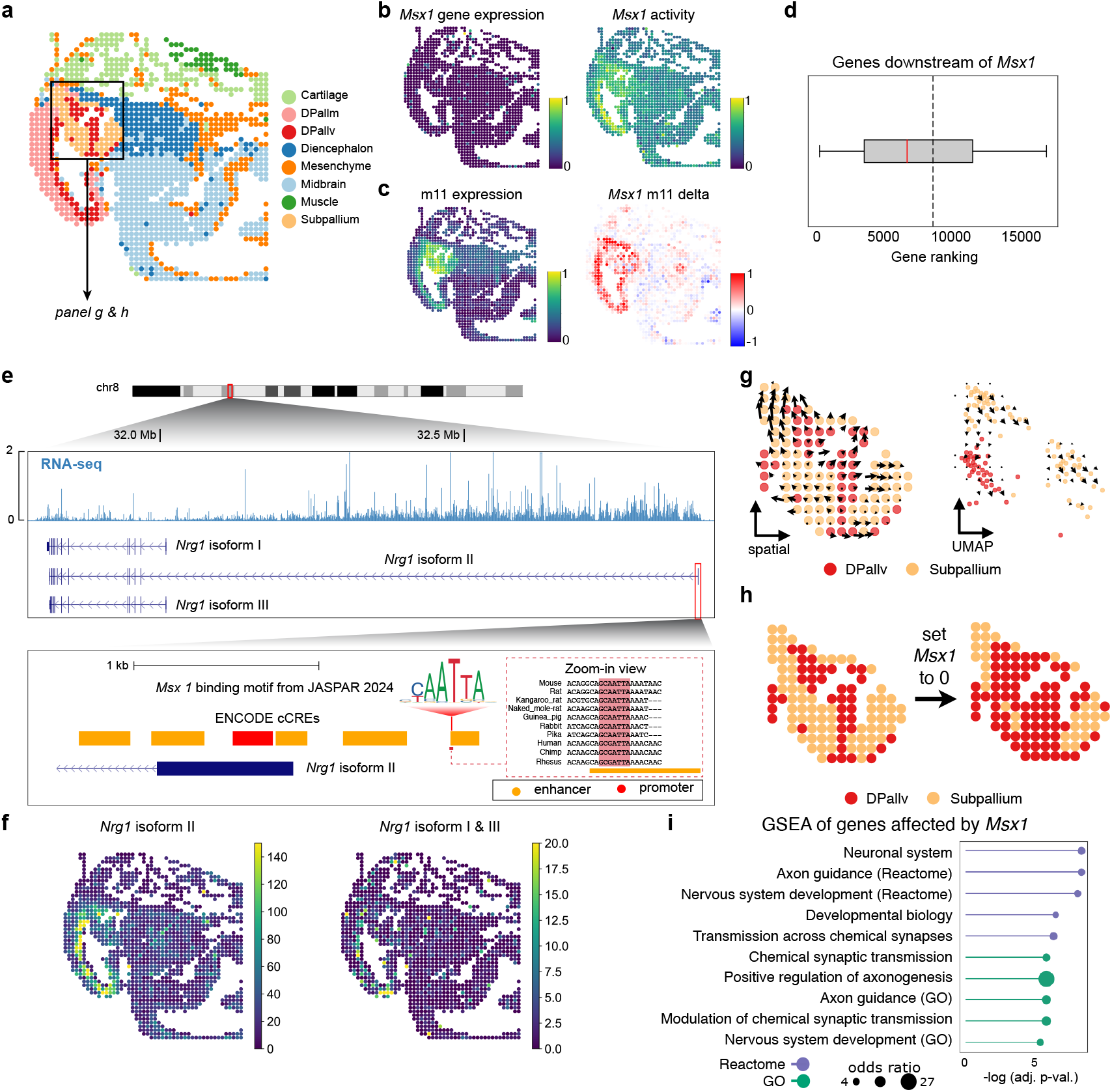
Eykthyr identifies *Msx1* as a key regulator of pallium differentiation. **a**. Spatial plot of a mouse embryo sample at E15.5, with spots annotated using Eykthyr. **b**. Spatial plots comparing sparse gene expression (left) of *Msx1* with denser TF activity (right) derived from chromatin accessibility data. **c**. Expression of metagene m11 (left) and predicted change in m11 expression following a simulated *Msx1* knockout (right). **d**. Box plot showing the ranking of genes downstream of *Msx1* from a generic mouse gene regulatory network [14]. **e**. Genomic view of *Nrg1* showing common splice variants, with isoform II containing large introns. The upper track shows RNA-seq read coverage from the E15.5 mouse embryo. The zoomed-in section highlights a *Msx1* binding site within an annotated proximal enhancer region near the transcription start site of *Nrg1* isoform II. **f**. Spatial expression of *Nrg1* isoform II (left) compared to all other isoforms (right). **g**. Visualizations of cell identity shifts following an *in silico* knockout of *Msx1*, showing transition from Subpallium to DPallv in both spatial (left) and UMAP (right) embeddings. **h**. Predicted region-specific identity shifts before and after simulated *Msx1* knockout. **i**. Gene set enrichment analysis of genes most affected by *Msx1*, revealing pathways associated with pallium development.

Notably, Eykthyr identified *Nrg1* as a gene strongly influenced by *Msx1*, despite not being previously annotated as a downstream target in existing GRNs. *Nrg1* spans more than 1 Mb and produces multiple isoforms (**Fig**. 3e). *Nrg1* isoform II includes an alternative transcription start site (TSS) located just 650 bp apart from a *Msx1* binding motif in an ENCODE [23] proximal cCRE region (**Fig**. 3e). This isoform has been linked to schizophrenia [24], a disorder recently associated with glutamatergic neurons in the dorsal-ventral axis [25]. *Nrg1* isoform II shows higher expression near the pallium at embryonic day 15 in mice [26], suggesting *Msx1* may directly regulate its activity during development. Its expression is largely confined to DPallv (**Fig**. 3f), indicating a potential role in regional differentiation. Eykthyr thus highlights a previously uncharacterized regulatory mechanism of *Nrg1* isoform II.

To further examine *Msx1*’s role, we performed *in silico Msx1* knockout and analyzed shifts in cell identity across spatial and metagene embeddings (**Fig**. 3g, h). Cells in the Subpallium shifted toward a DPallv-like identity, indicating *Msx1* plays a role in maintaining spatially distinct cellular fates. Gene set enrichment analysis of genes altered by the knockout confirmed *Msx1*’s role in neural development and axon guidance (**Fig**. 3i), including L1CAM upregulation [27, 28]. These changes were spatially specific, with minimal effects in other regions after the knockout.

This case study of *Msx1* highlights four key strengths of Eykthyr: (1) it detects low-expression TFs through chromatin accessibility, identifying regulators that might otherwise be overlooked; (2) it identifies region-specific regulatory effects; (3) its *in silico* knockout predictions aligh with known developmental roles; and (4) it captures subtle yet spatially impactful regulatory mechanisms.

### Identifying key regulators of radial glia differentiation

We next investigated a spatial ATAC–RNA-seq sample of the mouse embryonic hindbrain [8] (**Fig**. 4a and **Fig**. 5a), where radial glia differentiate into postmitotic premature neurons. Unlike the previous system, these cells follow a more continuous developmental trajectory. This setting allowed us to assess Eykthyr’s predictions under a different evaluation strategy based on pseudotime rather than discrete cluster separation. This trajectory is well suited to a spatial pseudotime approach, as postmitotic neurons spatially drift away from radial glia after differentiation, and the original study showed that spatial distance closely correlates with chromatin accessibility–based pseudotime.

**Figure 4:**
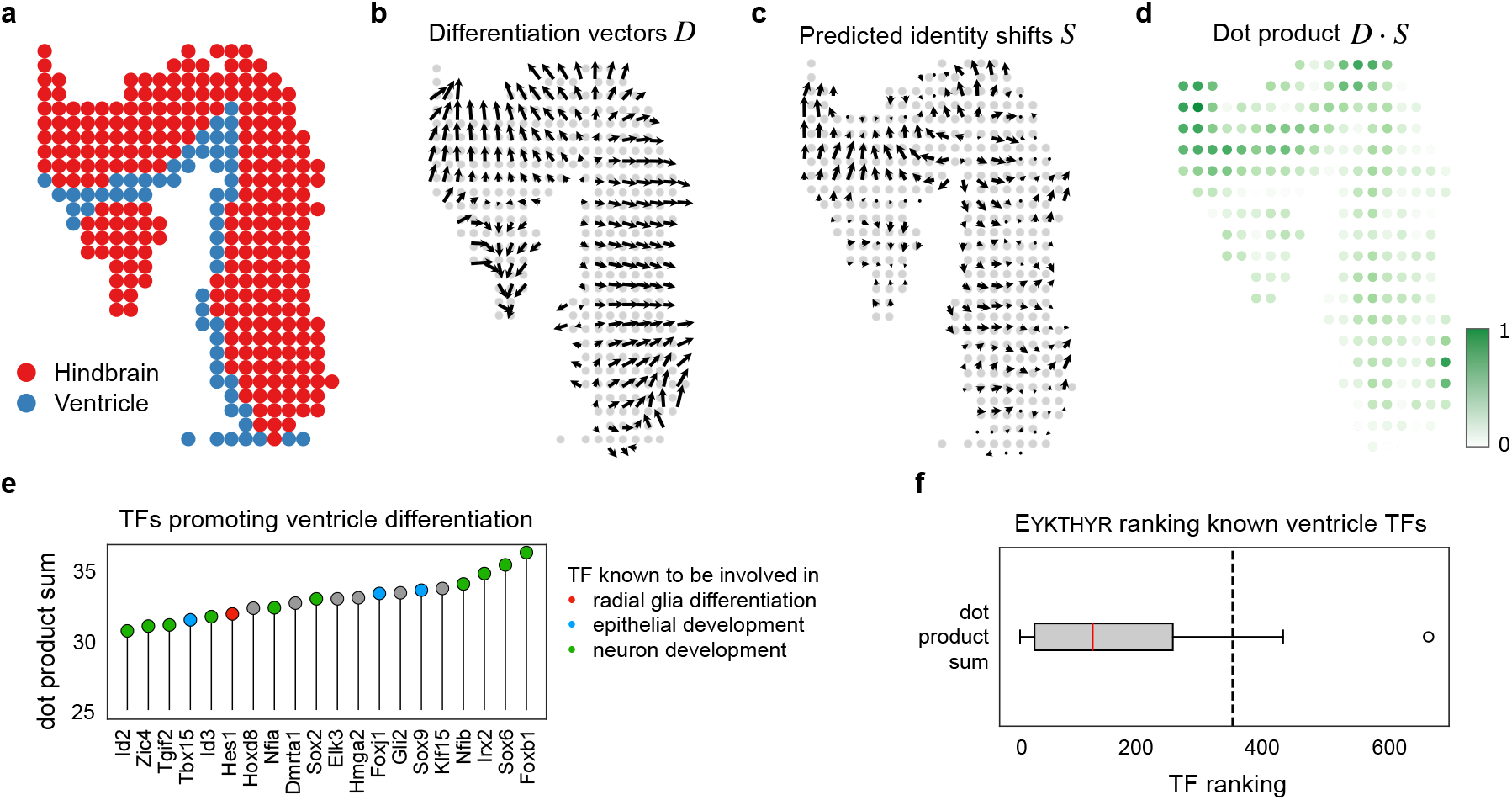
Evaluation of Eykthyr’s predictions for TFs driving radial glia differentiation. **a**. Spatial plot of a mouse embryo section from the spatial multiome (scATAC and scRNA-seq) dataset (further analyzed in **Fig**. 5). **b-d**. Metrics used to quantify a TF’s influence on promoting radial glia differentiation in the ventricle into postmitotic premature neurons in the hindbrain. The metric evaluates the alignment between the shift in cell identity due to (**b**) natural differentiation and (**c**) Eykthyr’s predicted metagene shift, by calculating (**d**) the dot product between these two vectors. **e**. Top TFs identified by Eykthyr as promoting differentiation from radial glia to postmitotic premature neurons, ranked according to this custom metric. **f**. Box plot displaying Eykthyr’s ranking of TFs known from the literature to influence this differentiation process. The dashed vertical line represents the expected median rank of a random set of TFs.

**Figure 5:**
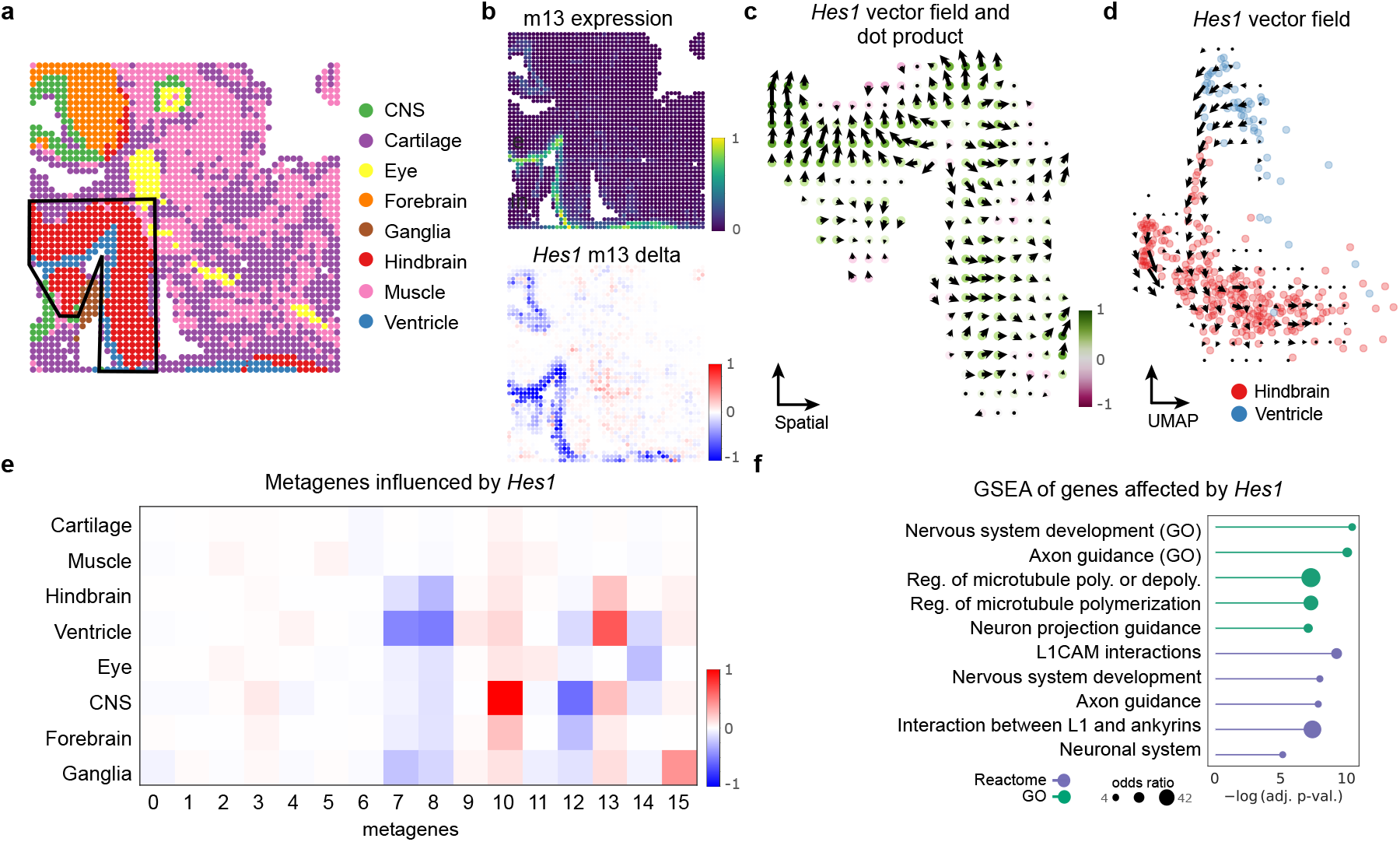
Eykthyr identifies *Hes1* as a key regulator in the differentiation of radial glia into postmitotic premature neurons. **a**. Spatial plot of a mouse embryo sample with spots annotated using Eykthyr. **b**. Expression of metagene m13 (top) and the predicted change in m13 expression following a simulated *Hes1* knockout (bottom). **c**. Eykthyr predicts that reducing *Hes1* activity would induce a shift in cell identity following the spatial trajectory of differentiation, underscoring its potential role in the differentiation process. **d**. UMAP visualization showing the transition in cell identity from Ventricle to Hindbrain after a simulated reduction in *Hes1* activity. **e**. Heatmap of metagenes most influenced by *Hes1* activity, highlighting regions of highest impact. **f**. Gene set enrichment analysis of genes most affected by *Hes1* in the Ventricle, revealing terms and pathways related to radial glia differentiation into postmitotic premature neurons.

We defined pseudotime as the normalized Euclidean distance of each spot to the nearest spot annotated as ventricle (**Fig**. 4b), approximating its progression along the radial glia-to-neuron differentiation path. After simulating Eykthyr’s TF knockouts (**Fig**. 4c), we computed directional vectors capturing how each cell’s metagene embedding shifts relative to its spatial neighbors. By comparing these vectors to the pseudotime axis using their inner product, we ranked TFs by how strongly their knockouts perturb the differentiation trajectory (**Fig**. 4d). We compiled a list of TFs from the literature associated with the differentiation of radial glia to postmitotic premature neurons [29], including *Ascl1, Dlx1, Dlx2, Emx1, Etv1, Gsx1, Gsx2, Hes1, Neurog2, Olig2, Pax6*, and *Sp8*.

Eykthyr successfully ranked these TFs with established roles in glial differentiation significantly higher than expected by chance (*P* = 0.02) (**Fig**. 4f). In contrast, the baselines – CellOracle and the expression-only variant of Eykthyr– showed weaker concordance with known regulators. Fourteen of Eykthyr’s top twenty ranked TFs included genes involved in ventricle differentiation (*Hes1*), neuron development (*Sox9, Foxj1*, and *Tbx15*), or epithelial development (*Foxb1, Sox6, Irx2, Nfib, Sox2, Nfia, Id3, Tgif2, Zic4*, and *Id2*) (**Fig**. 4e). These results reinforce that Eykthyr’s *in silico* knockout predictions are biologically grounded and accurately capture TFs responsible for spatially informed differentiation processes in the hindbrain and ventricle.

### *Hes1* as a spatial architect in radial glia differentiation

We performed a deeper analysis of Eykthyr’s predictions regarding *Hes1* and its role in the differentiation of radial glia into postmitotic premature neurons (**Fig**. 5a). While the original study emphasized factors like *Sox2* as key regulators of this process – which Eykthyr also identified – it did not highlight *Hes1*, which Eykthyr ranked highly (**Fig**. 4e).

Using Eykthyr’s framework, we identified metagene m13 as highly expressed in ventricle-proximal cells, with *Hes1* exerting strong regulatory influence on m13 in these early-stage populations (**Fig**. 5b). When *Hes1* activity was reduced, ventricle cells became more similar to hindbrain cells (**Fig**. 5c, d), recapitulating the known *Hes1*^-/-^ phenotypes of premature neuron production and depletion of progenitor cells [30].

Beyond the ventricle-hindbrain transition, Eykthyr predicted that *Hes1* regulates a different metagene (m10) in the central nervous system, suggesting that its regulatory role varies across tissue compartments (**Fig**. 5e). Leveraging the linear metagene embedding, we transformed the shifts in metagene expression back to gene-level changes and ranked genes by their predicted influence to *Hes1* perturbation. Comparing this ranked list to an orthogonal mouse GRN [14], we found significant enrichment of known downstream targets among the top-ranked genes (*P* = 8.1 × 10^−37^) (**Fig**. S3). Gene set enrichment analysis of these genes further revealed pathways related to glial differentiation, axon guidance, and nervous system development (**Fig**. 5f). Notably, L1CAM-associated processes were enriched, consistent with its established role in neuronal outgrowth and neurodevelopment [31].

These findings suggest that *Hes1* exerts spatially distinct regulatory effects along the ventricle-hindbrain axis, complementing previous findings on *Sox2*. More broadly, this analysis demonstrates how Eykthyr can reveal dynamic, region-specific roles of TFs in shaping developmental trajectories.

### Context-dependent regulation of T-cell states in tumor microenvironments

We applied Eykthyr to a single-cell Slide-tags multiome dataset of human metastatic melanoma [9], where two tumor compartments were defined based on distinct transcriptomic signatures in tumor cells (**Fig**. 6a). While the original study examined T-cell infiltration into each compartment using T-cell receptor (TCR) sequences, it did not focus on finer-grained T-cell subpopulations. Building on this, our metagene analysis revealed a proliferating T-cell population that was not clearly distinguished in the original clustering (**Fig**. 6b). Identifying this subset allowed us to assess how proliferating T cells respond differently across tumor microenvironment, rather than treating T cells as a single coarse population.

**Figure 6:**
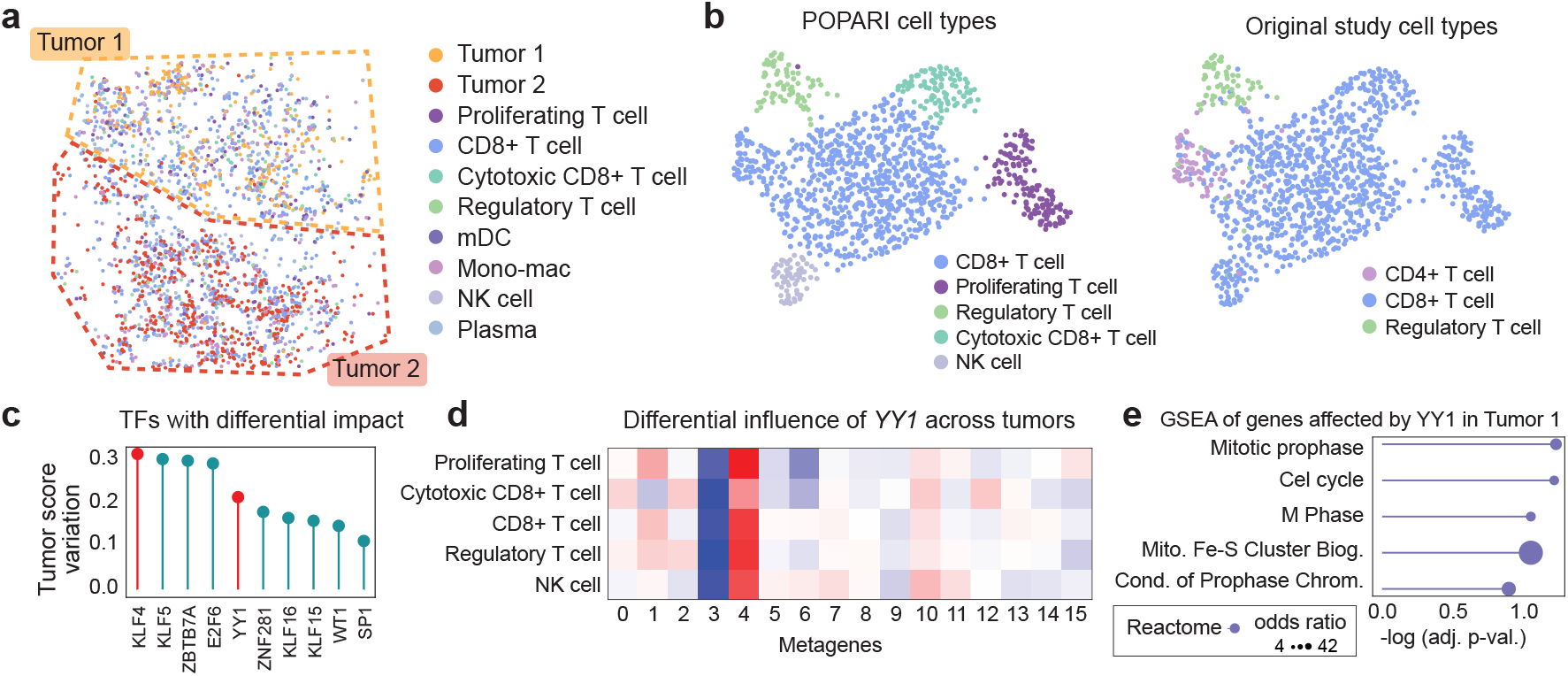
Eykthyr predicts a context-specific influence of *YY1* on T-cell states within distinct tumor microenvironments. **a**. Spatial plot of a human metastatic melanoma sample, with cells annotated by Eykthyr, showing spatial separation into two distinct tumor compartments. **b**. Comparison of T-cell annotations from the original study and those derived from Eykthyr’s embedding, revealing additional T-cell states identified by Eykthyr. **c**. Top TFs identified by Eykthyr ranked by their differential influence on Proliferating T Cells between the two tumor compartments. **d**. Heatmap displaying the context-specific influence of *YY1* on metagene expression across T-cell states, as measured by the difference in *YY1* influence between tumor compartments. **e**. Gene set enrichment analysis of genes with both a *YY1* binding motif in their promoter and a cross-tumor difference in promoter accessibility in Proliferating T Cells.

To uncover the transcriptional drivers of proliferating T cells, we used Eykthyr’s *in silico* knockout framework. This analysis identified *KLF4* and *YY1* as key regulators with compartment-specific effects on proliferating T cells (**Fig**. 6c). *KLF4* has been linked to tumor suppression and improved survival in human cancers [32], and it can either reverse CD8 T-cell exhaustion or limit T-cell proliferation, depending on context [33]. Eykthyr’s predictions reflected this duality: in one compartment, *KLF4* knockout increased predicted T-cell proliferative potential, while in the other, it appeared to have an inhibitory effect.

Similarly, *YY1* has been shown to promote or inhibit T-cell proliferation in a context-dependent manner [34, 35]. Analyzing transitions in T cell state before and after an *in silico YY1* knockout revealed distinct shifts in metagene expression – particularly among m3 and m4 – across compartments (**Fig**. 6d). These shifts correlated with differential chromatin accessibility at *YY1* binding sites, suggesting that *YY1* may alter proliferating T-cell function in a compartment-specific manner. To explore this further, we collected genes with a *YY1* motif in their promoter region and assessed their chromatin accessibility between tumor compartments. Gene set enrichment analysis on genes with higher accessibility in the bottom tumor compartment showed enrichment for cell-cycle pathways (**Fig**. 6e), whereas no significant cell-cycle enrichment was observed in the top compartment, indicating enhanced proliferation from *YY1* only in the bottom compartment.

These findings illustrate how Eykthyr identifies TFs that regulate T-cell states in a compartment-specific manner, shaping tumor-immune interactions. Understanding how *KLF4* and *YY1* drive or restrain T-cell proliferation in distinct microenvironments may inform further immunotherapeutic strategies.

Note that to further test the robustness of Eykthyr’s GRN design, We also evaluated a modified, non-spatial version of Eykthyr on a zebrafish scRNA-seq dataset with *in vivo* knockout experiments. Results are provided in the **Supplementary Note**. Together, these analyses highlight how Eykthyr’s *in silico* knockout simulations can reveal biologically meaningful regulators in both spatial and non-spatial settings.

## Discussion

We introduced Eykthyr, a computational framework that integrates spatial transcriptomic and chromatin accessibility data to identify location-specific regulators of gene programs in complex tissues. By inferring TF activity directly from chromatin accessibility, Eykthyr overcomes high dropout and low transcript signals that often obscure key regulators. Across diverse applications – including embryonic brain development and compartment-specific T-cell states in melanoma – Eykthyr identified TFs with minimal gene expression but strong regulatory influence. Its *in silico* TF knockouts further reveal which cells or regions are most affected by a given TF perturbation, underscoring the spatial dependence of gene regulation. Compared to existing methods that rely on single-modality or non-spatial data, Eyk-thyr is the first framework explicitly designed to leverage both spatial transcriptomics and chromatin accessibility, offering new insights into how local environments shape cell identity.

Despite its strengths, Eykthyr has several limitations. The required paired spatial multiome data – an emerging but still not widely available resource – and our validations have focused on relatively discrete developmental and tumor contexts. Future work should expand to tissues with overlapping lineages or higher cellular complexity, such as adult organs or advanced disease states. Additionally, Eykthyr assumes a fixed spatial map, limiting its ability to simulate large-scale morphological changes such as compartment merging or regional emergence. Addressing these challenges will require more flexible spatial models and expanded datasets that better capture biological complexity.

Several extensions could enhance Eykthyr’s modeling capabilities. Currently, it assumes that metagene expression is a linear combination of TF activity. Yet transcriptional regulation is often non-linear: TFs can act cooperatively or exhibit threshold effects, and chromatin context or epigenetic states can modulate TF activity in complex, context-dependent ways that linear models cannot fully capture. Future version of Eykthyr could incorporate spatio-temporal modeling to analyze tissue samples collected over time. This would allow dynamic models to capture lagged TF effects, identifying how early regulatory signals shape later gene expression. Joint modeling across multiple samples or developmental stages could further improve robustness and reveal consistent regulatory trajectories. Adapting Eyk-thyr to unpaired spatial transcriptomic and single-cell multiomic data represents another important direction, broadening applicability to more datasets. Integrating additional modalities, such as proteomics, or shifting from spot-level to true single-cell resolution may refine TF activity estimates, as each data type captures only part of a TF’s regulatory influence.

Beyond model enhancement, Eykthyr’s *in silico* TF perturbation framework offers a valuable tool for hypothesis generation. By predicting how TF activity shifts reshape tissue architecture and cell state transition, the model can help prioritize candidate regulators for targeted experimental validation. Methods such as PerturbView [36], Perturb-FISH [37], and Perturb-map [38] combine *in situ* molecular profiling with multiplex antibody staining to measure spatial perturbation effects [39]. Integrating Eyk-thyr’s predictions with these techniques could accelerate the discovery of key regulatory circuits and support iterative spatial perturbation experiments to uncover causal mechanisms of tissue organization.

Overall, by pinpointing region-specific TF effects, Eykthyr provides a solid foundation for precision spatial gene regulatory studies and paves the way for targeted strategies to modulate TF activity for a wide range of tissue contexts in health and disease.

## Methods

### The Eykthyr model

Eykthyr takes spatial multiome data as input and generates low-dimensional embeddings for both gene expression (as metagenes) and chromatin accessibility (as TF activity) for each cell or spot. Each embedding dimension is a linear combination of the input data, preserving interpretability – shifts in the embedding directly reflect changes in gene expression or chromatin accessibility. Eykthyr then learns the relationship between these embedding spaces for each cell by training a linear model based on the embeddings of spatially proximal cells within the tissue. It can then simulate how changes in TF activity would affect downstream metagene expression using these learned linear relationships.

#### Constructing linear embedding space from spatial transcriptome data

Eykthyr addresses spatial transcriptomic sparsity by generating low-dimensional metagene embeddings using Popari [17]. Popari takes preprocessed read counts (filtered, normalized, and log transformed with SCANPY [40]) and incorporates spatial information via a hidden Markov random field framework. In short, Popari identifies *K* distinct metagenes that capture the fundamental transcriptomic factors within one or more samples. A metagene is defined as a latent factor representing a combinatorial signature of coexpressed genes with spatial coherence. These factors are encoded in a *G* × *K* metagene matrix, **D**, where *G* is the total number of genes measured. Each cell is represented by a *K*-dimensional latent vector, **M**_*i*_, which describes its state in the metagene space, and its observed gene expression **Y**_*i*_ is modeled as a nonnegative linear combination of the metagenes:

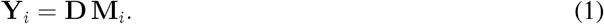

Popari also estimates a *K* × *K* spatial affinity matrix **Λ** for each sample, quantifying the coexpression of metagene pairs among neighboring cells. The metagene matrix **D**, the latent embeddings **M**_*i*_, and the spatial affinity matrices **Λ** are jointly optimized via an alternating maximum *a posteriori* (MAP) procedure, yielding intuitive representations of latent spatial structure. Each column **d**_*k*_ of **D** provides an interpretable ranked list of genes, with the top-ranked genes forming a spatially coherent module. This approach enables the capture of nuanced combinatorial effects within a denoised, low-dimensional embedding, facilitating spatially-aware interactions between TFs and downstream gene expression even in high-noise or sparse settings.

#### Estimating transcription factor activity from spatial epigenome data

To address limited expression of TFs, Eykthyr estimates TF activity directly from chromatin accessibility data using ArchR [41] and following the MISAR-seq data processing workflow [7]. Specifically, we generate Arrow files with 5kb-tile matrices and call peaks using MACS2 [42]. We then apply a latent semantic indexing (LSI) transformation to the most accessible peaks, followed by shared nearest neighbor clustering to form initial clusters. These clusters are then refined using spatial cell location information: cells with a different cell type than all of their neighbors are are reassigned based on the most enriched cluster among their *k* nearest neighbors.

We then recalculate the most variable peaks within these refined clusters and use these features for the next iteration of LSI. This process is repeated until cluster assignments converge. Finally, we annotate peaks with TF motifs using ArchR’s addMotifAnnotations function, which scans peak sequences for TF binding motifs from the CIS-BP database [43], retaining only motif matches with *p <* 1 × 10^−5^ to ensure statistically significance.

Given the cell-by-peak matrix **P** and the peak-by-TF-motif matrix **T**, TF activity estimates for each cell are computed as:

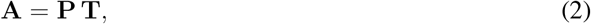

which gives the number of times each TF motif appears in a peak measured for a given cell. To account for motif prevalence, we normalize TF activity across all cells for each TF to the range [0, 1].

#### Determining TF influence on metagenes

Conceptually, TF influence is modeled as a bipartite graph between TFs and metagenes, with directed edges from each TF to each metagene, and edge weights representing the influence of a TF on metagene expression. To compute these edge weights, we use a regularized linear model, fit separately for each cell using the metagene expression and TF activity of its *n* spatially proximal neighbors. We choose *n* to include a sufficient number of cells for the model while avoiding those too distant. Since spatial multiome datasets vary in resolutions (e.g., spot-based vs. single-cell), *n* is dataset-specific, though we found *n* = 100 to be effective across datasets analyzed.

For each cell, we construct a neighborhood comprising spatially proximal cells – half from the same cell type, and half unrestricted. We train the model to predict the metagene expression matrix **M**_*i*_ for the neighborhood of cell *i* based on the TF activity matrix **A**_*i*_ and a learned weight matrix **W**_*i*_

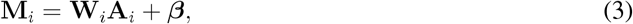

where ***β*** is a vector of biases (one per metagene). This formulation assumes potential regulatory influence of each TF on all metagenes. Although not all TF interactions are additive, it provides a simple model that is robust to high noise and sparsity inherent to spatial multiome data. Another advantage of this formulation is that it allows for interpretable simulations, as the relationship between gene expression, chromatin accessibility, metagene expression, and TF activity remains linear.

To avoid overfitting, we employ L2 regularization via Ridge regression [44] with regularization strength *α* = 1, and further enhance the robustness of the model by bagging. Specifically, we train 10 estimators on bootstrap samples, each with 80% of randomly selected features, and combine their output to obtain a posterior distribution over the coefficients,

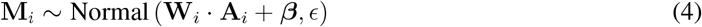

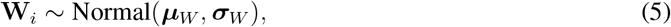

where ***µ***_*W*_ and ***σ***_*W*_ are matrices of shape (*k* × *t*)), with each ***µ***_*k,t*_ and ***σ***_*k,t*_ representing the mean and standard deviation for the effect of TF *t* on metagene *k*. This ensemble approach reduces variance by averaging the coefficients across multiple bootstrapped models, producing more robust estimate of TF influence on metagene expression.

#### Simulating changes in TF activity

After learning TF–metagene interactions, we simulate changes to TF activity to predict their impact on metagene expression. For each TF *j*, we set its activity to zero across all cells – simulating an experimental knockout – effectively removing its regulatory influence across the tissue. The adjusted metagene expression matrix for cell 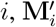,is computed as:

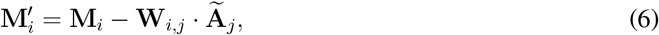

where 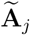 represents the change to **A** from setting TF *j*’s activity to zero.

By performing this tissue-wide *in silico* TF knockout, we aim to uncover how gene expression and cell identity might change in the absence of a specific TF. A TF knockout is more straightforward to simulate than upregulation or downregulation, as it is unclear how to realistically increase or decrease TF activity across all cells.

While Eykthyr’s simulated knockouts reliably indicate the direction of change in the metagene embedding space, the magnitude of change is often small due to strong regularization. Nevertheless, the relative ranking of TF effects remains robust for downstream tasks such as identifying developmental regulators. For downstream applications that require a more pronounced signal – such as visualizing cell identity changes – we apply a scaling factor to amplify magnitude shifts, enhancing clarity in visualizations and robustness in clustering analyses. For the visualizations of cell state transition (**Figs**. 3g, 3h, 4c, and 5d), we used a scaling factor of 10.

### Evaluating Eykthyr’s performance

To evaluate Eykthyr’s predictions of TF influence, we developed visualization techniques, statistical tests, and scoring metrics to assess shifts in cell embeddings following simulated TF perturbations.

#### Scoring known TFs using batch correction metrics

Validating Eykthyr’s predictions is challenging due to the limited availability of spatially-informed experimental TF knockouts. We leveraged the biological principle that cell types derived from common progenitors exhibit similar metagene expression patterns. Thus, increased similarity among these cell types after simulated TF knockout suggests a regulatory role for that TF in their differentiation.

To assess this, we adapted batch correction metrics – traditionally used to evaluate cell mixing across datasets – to evaluate TF perturbation effects. Using metrics from the scib-metrics Python package [45], including average silhouette width, graph iLISI, and principal component regression, we quantified the extent to which differentiated cells mix in the embedding space following simulated TF knockouts.

These metrics yield scores reflecting each TF’s predicted impact on differentiation. We ranked TFs based on these scores and evaluated the statistical significance of their rankings using a t-test, focusing on TFs previously linked to differentiation. This approach provides a statistical framework for validating Eykthyr’s biologically meaningful predictions.

#### Calculating pseudotime from spatial distance

Pseudotime provides an estimated measure of a cell’s progression through a developmental process and is commonly used to study differentiation. Each cell is assigned a value along a pseudotime axis, enabling analysis of gene regulation relative to differentiation trajectories. While pseudotime is typically computed from transcriptomic embeddings, many tissues exhibit spatial stability, where cells remain physically proximal to their progenitors, allowing spatial distance to serve as a proxy for pseudotime.

Given a set of progenitor cells *P* and differentiated cells *C*, we define the pseudotime *t*(*c*) of each cell *c* ∈ *C* as its minimum distance to any progenitor:

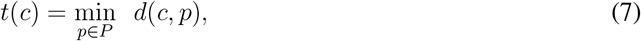

where *d*(*a, b*) is the Euclidean distance between *a* and *b*. This distance is normalized across all cells in *C* to the range [0, 1].

This spatially defined pseudotime serves as a directional vector of differentiation within the tissue. To evaluate TF effects, we examine each cell’s nearest spatial neighbors and identify the neighbor it most resembles after a simulated TF knockout. This yields two directional vectors per cell: one for the differentiation trajectory *d* and one for the TF knockout effect *s*. We compute the magnitude of their inner product, |⟨*d, s*⟩|, and rank TFs by their projected impact on the differentiation process.

#### Benchmarking against CellOracle

We benchmarked Eykthyr against CellOracle using the MISAR-seq spatial multiome dataset, focusing on the E15.5-S2 slice, as well as the spatial ATAC–RNA-seq mouse embryo dataset. CellOracle was run using its standard preprocessing steps outlined in its tutorial, with spatial information omitted, aligning with its design as a non-spatial method. We note that the dataset likely contains more than one cell per spot, which poses additional challenges for CellOracle, as it is optimized for single-cell data.

In preprocessing, we retained all highly variable genes and included all TFs used by Eykthyr for this dataset, excluding TFs without detected transcripts. CellOracle’s clustering was performed using its non-spatial method, producing clusters that were less refined than those generated by Popari but sufficient for this benchmark.

For GRN construction, we tested both CellOracle’s default mouse GRN (derived from a large scATAC-seq dataset) and a sample-specific GRN built per its guidelines. The sample-specific GRN underperformed, likely due to sparse chromatin accessibility data and limited cell numbers, so we used the default GRN for our comparisons.

To quantitatively compare performance, we applied the batch integration evaluation framework to the MISAR-seq dataset and the spatial pseudotime procedure to the spatial ATAC–RNA-seq dataset, as described above.

### Visualizations

To analyze the downstream effects of TF knockouts, we used visualization techniques inspired by CellOracle and adapted from RNA velocity approaches in the Velocyto package [46]. CellOracle, designed for non-spatial scRNA-seq data, computes cell-state transition probabilities based on changes in gene expression. To visualize cell identity changes after setting TF activity to zero, CellOracle requires both (i) the predicted gene expression change and (ii) the gene expression–based cell embedding.

For Eykthyr, we adapted this framework by replacing gene expression changes with changes in metagene expression. We generated a two-dimensional embedding of the metagenes using partition-based graph abstraction (PAGA) [47], which preserves the data’s topological structure. Both metagene embeddings and spatial cell locations were used as embedding spaces for downstream analysis. With these adaptations, we applied CellOracle’s transition probability framework to compute how each cell shifts relative to its neighbors in either spatial or metagene space. These probabilities were then converted into directional vectors and combined into a vector field using an averaging approach from Velocyto.

To visualize metagene affinities, we used Popari’s empirical correlation plotting functions. For assessing changes in Leiden clustering, we used Popari’s function to identify the optimal resolution for a specified number of clusters.

A detailed description of dataset-specific preprocessing, Eykthyr parameter selection, and analysis methods is provided in the **Supplementary Note**.

## Acknowledgment

This work was supported, in part, by National Institutes of Health Common Fund 4D Nucleome Program grant UM1HG011593 (J.M.); National Institutes of Health Common Fund Cellular Senescence Network Program grant UH3CA268202 (J.M.); and National Institutes of Health grants R01HG007352 (J.M.), R01HG012303 (J.M.), R21DA061481 (J.M.), and U24HG012070 (J.M.). J.M. was additionally supported by the Ray and Stephanie Lane Professorship, a Guggenheim Fellowship from the John Simon Guggenheim Memorial Foundation, a Google Research Award, and a Single-Cell Biology Data Insights award from the Chan Zuckerberg Initiative. S.K. is a Lane Fellow. The funders had no role in study design, data collection and analysis, decision to publish or preparation of the manuscript.

## Code Availability

The source code for Eykthyr can be accessed at: https://github.com/gkrieg/eykthyr.

## Data Availability

All datasets analyzed in this work are publicly available. The MISAR-seq mouse embryo dataset was obtained from ref. [7]; the whole mouse embryo dataset was obtained from ref. [8]; the human metastatic melanoma dataset was obtained from ref. [9] and the zebrafish scRNA-seq dataset was obtained from ref. [14].

## Author Contributions

Conceptualization, S.K., J.M.; Methodology, S.K., J.M; Software, S.K.; Investigation, S.K., E.H., J.M; Writing, S.K., J.M; Funding Acquisition, J.M.

