## Supplemental Information for "EYKTHYR reveals transcriptional regulators of spatial gene programs"

#### Supplementary Figures

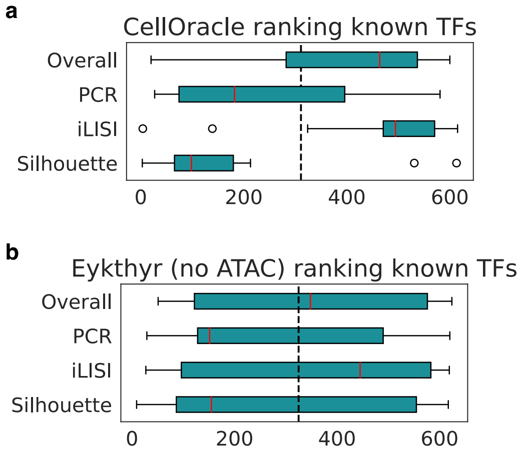


**Figure S1:** Extended validation on the MISAR-seq dataset. **a.** Box plot showing CellOracle's ranking of TFs associated with pallium differentiation (*Arx*, *Emx1*, *Emx2*, *Gli3*, *Lef1*, *Lhx2*, *Lmx1a*, *Nr2e1*, *Nr2f1*, *Nr2f2*, *Otx1*, *Bhlhe22*, *Pax6*, and *Sp8*). The overall score shows that CellOracle predicts these TFs have an inhibitory effect on differentiation, and is nearly statistically significant ($P = 0.07$). **b.** Box plot showing EYKTHYR's ranking of the same TFs. Instead of using the chromatin accessibility information, EYKTHYR was only allowed to use the gene expression of TFs to estimate TF activity. Overall, EYKTHYR's predictions are essentially random when no chromatin accessibility is used.

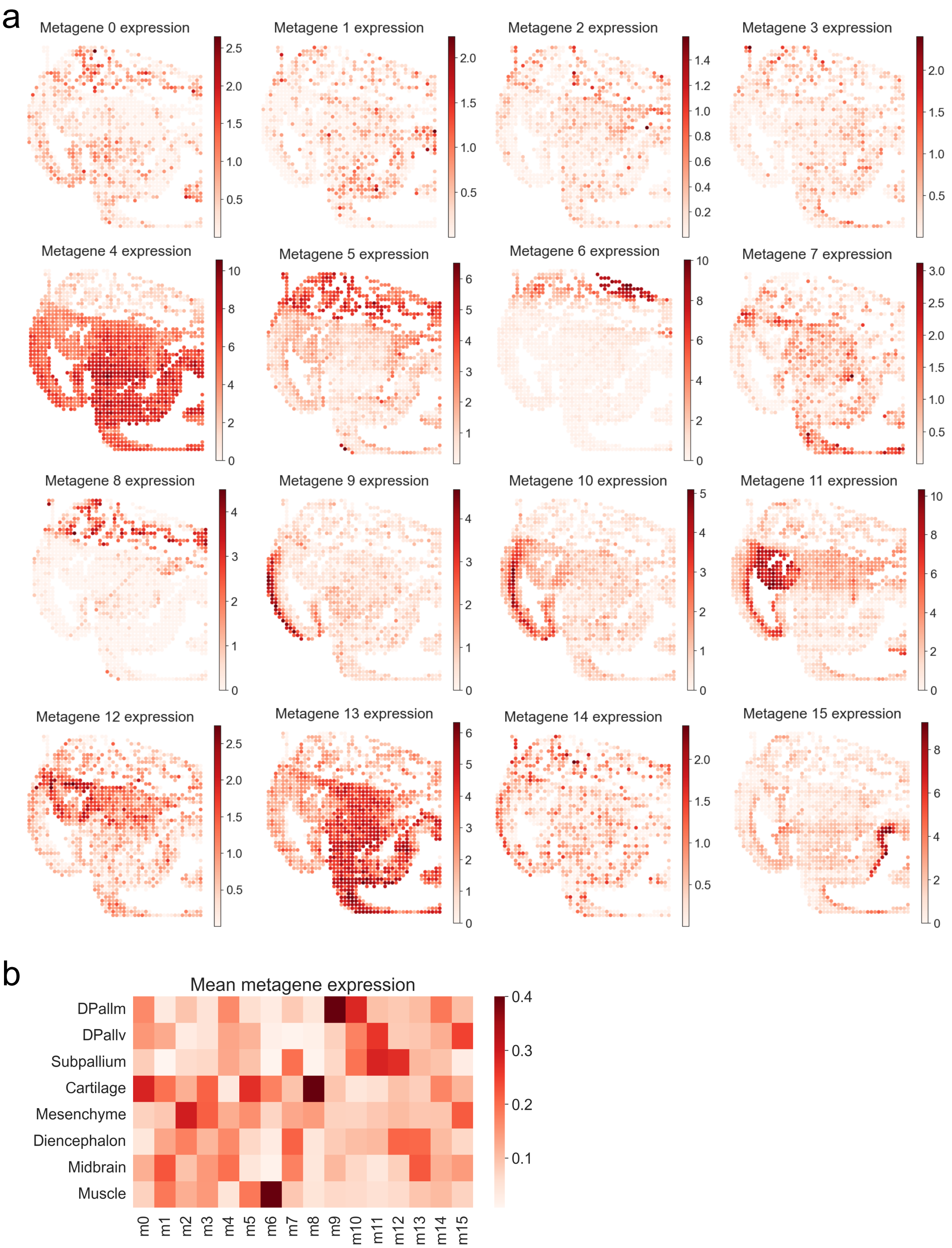


**Figure S2:** Extended metagene analysis for the MISAR-seq dataset. **a**. *In situ* metagene plots for the MISAR-seq dataset, slice E15_5-S1. **b**. Heatmap of mean expression of each metagene by each cell type, normalized by column.

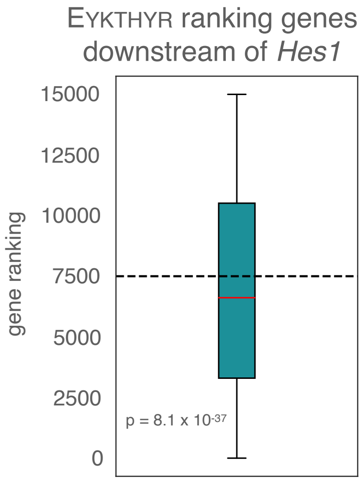


**Figure S3:** Analysis of genes influenced by *Hes1* from the spatial ATAC–RNA-seq dataset. Box plot showing the ranking of genes downstream of *Hes1* within CellOracle's default mouse GRN. The ranking is given by EYKTHYR's predicted influence of *Hes1* on downstream genes.

**a**

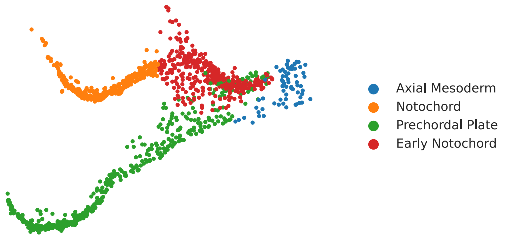


**b**

*Sebox* knockout

| Cell type | Early axial mesoderm | Early notochord | Notochord | Prechordal plate |
|---|---|---|---|---|
| Experimental abundance shift | ↑ | ↑ | ↓ | ↓ |
| Eykthyr predicted shift | ↑ | ↑ | ↓ | ↓ |
| CellOracle predicted shift | ↑ | ↑ | ↓ | ↓ |

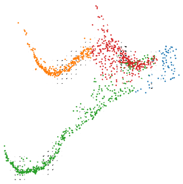

**c**

*Lhx1a* knockout

| Cell type | Early axial mesoderm | Early notochord | Notochord | Prechordal plate |
|---|---|---|---|---|
| Experimental abundance shift | ↑ | ↑ | ↓ | ↓ |
| Eykthyr predicted shift | ↑ | ↑ | ↓ | ↓ |
| CellOracle predicted shift | ↑ | ↑ | ↓ | ↓ |

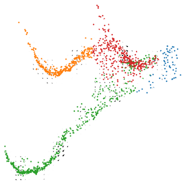

**d**

*Irx3a* knockout

| Cell type | Early axial mesoderm | Early notochord | Notochord | Prechordal plate |
|---|---|---|---|---|
| Experimental abundance shift | – | – | – | – |
| Eykthyr predicted shift | – | – | – | – |
| CellOracle predicted shift | ↑ | ↑ | ↓ | ↓ |

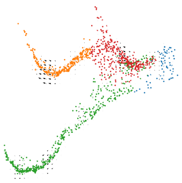

**Figure S4:** Results of experimental and *in silico* knockouts in scRNA-seq zebrafish dataset. **a**. Embedding of cells before knockout, colored by their original cell type. **b**. Table showing the change in the abundance of each cell type following a *Sebox* knockout. Ground truth shift in abundance is shown alongside the predicted shift of both EYKTHYR and CellOracle. **c**. Table showing the change in the abundance of each cell type following a *Lhx1a* knockout. Ground truth shift in abundance is shown alongside the predicted shift of both EYKTHYR and CellOracle. **d**. Table showing the change in the abundance of each cell type following an *Irx3a* knockout. Ground truth shift in abundance is shown alongside the predicted shift of both EYKTHYR and CellOracle.

### Supplementary Note

#### Computing metagenes with POPARI

We computed spatial metagenes using POPARI [1] in a two-phase optimization (first without, then with spatial-affinity updates) for each dataset. For the MISAR-seq mouse embryonic development slices (E11.0-S1, E13.5-S1, E15.5-S2, E18.5-S1), we set the number of metagenes $K = 16$, the spatial affinity regularization `lambda_Sigma_x_inv` $= 10^{-5}$, the inter-slice regularization `lambda_Sigma_bar` $= 10^{-2}$, and used "differential lookup" mode; after preprocessing we ran 10 iterations without spatial-affinity optimization, then 200 iterations while optimizing those parameters. For the single-sample Spatial ATAC–RNA-seq mouse embryo dataset [2], which exhibits large coherent regions, we omitted binning and inter-slice affinities, used $K = 16$ with `lambda_Sigma_x_inv` $= 10^{-4}$, then the same 10/200 iteration schedule. In the Slide-tags multiome metastatic melanoma dataset [3], due to single-cell resolution and tissue heterogeneity we ran POPARI in hierarchical mode (default binning) with $K = 16$ and `lambda_Sigma_x_inv` $= 10^{-4}$, performed the 10/200 iterations, and finally applied the superresolution function for 200 iterations to project embeddings to single-cell resolution.

#### Computing average metagene change after knockout

To quantify how each *in silico* perturbation shifts metagene activity, we computed, within each cell type, the average per-cell change in metagene expression and displayed the results as a heatmap using the "bwr" colormap centered at zero. In the MISAR-seq dataset, we measured this change following an *Msx1* knockout; in Spatial ATAC–RNA-seq, we applied the same procedure to a *Hes1* knockout; and in the Slide-tags multiome data we averaged metagene changes after a *YY1* knockout separately for each tumor compartment.

#### Visualization of cell-identity shifts

We projected knockout-induced cell-identity shifts as arrows on both spatial maps and PAGA graphs, drawing on Velocyto's styling [4] and CellOracle's strategy [5]. For the MISAR-seq and Spatial ATAC–RNA-seq datasets, we first ran a random-noise control to confirm that no arrows appear spuriously, then tuned arrow-length, width, and scale until shifts were clearly visible without clutter.

#### Gene set enrichment analysis

We transformed metagene-level perturbations into gene-level changes by multiplying the average metagene change vector $\Delta m$ by the metagene matrix $M$, yielding $\Delta g = M\,\Delta m$. After ranking genes by $\Delta g$ within each cell type, we selected the top 200 genes in those cell types whose count of nonzero hits exceeded 200/#celltypes. We then ran GSEApy [6] against the `GO_Biological_Process_2023` and `Reactome_2022` collections, reporting the top five terms per database and using the union of both gene sets as background. In MISAR-seq, only DPallv, DPallm, and Subpallium passed enrichment; in Spatial ATAC–RNA-seq, Ventricle, Hindbrain, and CNS (with Ventricle dominant); and in the Slide-tags multiome data we defined the universe $G$ as all genes with a *YY1* motif within 10 kb of the TSS, scored each by compartment-normalized ATAC fragment counts, ranked the top 200 differential genes (top vs. bottom compartment and vice versa), and found significant Reactome pathways (no GO terms survived FDR).

#### Differential transcription factor analysis

To identify TFs driving T-cell differentiation across tumor regions in the Slidetags-multiome dataset, we applied scib-metrics [7] to score each TF's effect on proliferating and CD8+ cytotoxic T cells within each compartment. We then ranked TFs by the absolute difference between compartment-specific scores, highlighting those with the greatest differential impact on T-cell fate.

#### Validation on zebrafish scRNA-seq data (direct comparison to CellOracle)

Due to the sparsity of spatial multiome data, EYKTHYR estimates the influence TFs have on metagenes, instead of on gene expression directly. To assess this design choice, we compared EYKTHYR's predictions to CellOracle's simulations using an scRNA-seq dataset with four experimental TF knockouts from zebrafish, as detailed in their original paper [5].

For this comparison, we adapted EYKTHYR to handle non-spatial, non-multiome data. First, given the lack of chromatin accessibility data, we used TF gene expression levels to estimate TF activity, which was feasible due to the reduced sparsity in this dataset. Second, because spatial information was not available, we calculated a single set of edge weights between TFs and metagenes for each cell type, mirroring CellOracle's approach to GRN construction.

For each TF, we simulated setting its activity to zero, and compared the resulting changes in cell type proportions to those observed in the corresponding experimental knockout sample (where TF activity was indeed set to zero). To compute changes in cell type proportions with EYKTHYR, we first calculated cell identity transition vectors for each cell. We then conducted a Markov random walk simulation, following the same methodology used by CellOracle and utilizing the differentiation vectors provided by CellOracle for this dataset.

The wildtype zebrafish dataset from Farrell et al. [8] served as the control for simulations, while the *Tyr* knockout dataset from Kamimoto et al. [5] served as the experimental control for TF knockouts. To compare the cell identity shifts calculated by EYKTHYR with both the experimental knockouts, and CellOracle, we calculated the proportional change in cell type identity in the experimental knockouts and normalized it to the control used in the simulation.

CellOracle's complete GRN model correctly predicted phenotypic changes for only three of four tested TF knockouts (**Fig.** S4b–d), whereas EYKTHYR successfully captured all four. These findings suggest that EYKTHYR's simplified GRN better models shifts in cell identity, even without spatial or accessibility data, reinforcing the biological validity of its *in silico* knockouts.
